# Inclusion of database outgroups improves accuracy of fungal meta-amplicon taxonomic assignments

**DOI:** 10.1101/2022.11.21.517387

**Authors:** Clayton Rawson, Geoffrey Zahn

## Abstract

Meta-amplicon studies of fungal communities rely on curated databases for assigning taxonomy. Any host or other non-fungal environmental sequences that are amplified during PCR are inherently assigned taxonomy by these same databases, possibly leading to ambiguous non-fungal amplicons being assigned to fungal taxa. Here, we investigated the effects of including non-fungal outgroups in a fungal taxonomic database to aid in detecting and removing these non-target amplicons. We processed 15 publicly available fungal meta-amplicon data sets and discovered that roughly 40% of the reads from these studies were not fungal, though they were assigned as *Fungus sp*. when using a database without non-fungal outgroups. We discuss implications for meta-amplicon studies and recommend assigning taxonomy using a database with outgroups to better detect these non-fungal amplicons.

## INTRODUCTION

Mycobiome studies commonly use high-throughput sequencing of either the ITS1 or ITS2 region amplified from environmental DNA (meta-amplicons) to investigate fungal diversity and community structure. These ITS barcode regions are useful for identifying fungal species, but are hypervariable and thus unsuitable for alignment and phylogenetic placement. In order to assign taxonomy to ITS amplicons, the reads must be compared against a database of known fungi. Thus, fungal meta-amplicon studies are limited by the databases available to match unknown amplicons against. Popular fungal sequence databases such as UNITE (Abarenkov et al., 2010), ISHAM (Irinyi et al., 2015), and Maarjam (Öpik et al., 2010), have been developed and maintained by the mycological community to serve as improvements over uncurated databases such as the NCBI non-redundant nucleotide database. These databases benefit from expert curation and are useful for querying unknown fungal sequences, but they may not be suitable for assigning taxonomy to meta-amplicon sequences due to their lack of outgroups.

Common fungal primers for the ITS1 and ITS2 regions can co-amplify other taxa from the environment, such as plants and metazoans (Bellemain et al., 2010). Despite what *in silico* tests might suggest, real-world PCR conditions can allow more permissive binding to non-target templates than might be expected. These non-target taxa that amplify will subsequently be sequenced and their amplicon reads will be returned among the fungal reads of interest. These reads must be removed before any downstream quantification of fungal diversity or community structure can proceed, but detecting them can prove to be tricky when using common bioinformatic pipelines.

The danger lies in using a curated database containing only fungi when assigning taxonomy to unknown amplicons. Without outgroups present in the database, non-fungal reads can be assigned to “Unknown Fungus.” Taxonomic assignment methods such as the Ribosomal Database Project Classifier (Wang et al., 2007), Least Common Ancestor (Hanson et al., 2016), or simply using the top BLAST (Altschul et al., 1990) hit will all attempt to find the closest match to a query sequence in a given database. If a reasonable match cannot be made at a given taxonomic level, the assignment algorithms find the most inclusive level at which taxonomy can be assigned. This leads to assignments such as k Fungi; p Ascomycota; c Sordariomycetes; o Xylariales; f Xylariaceae; g NA; s NA. In this example, taxonomy could only be assigned unambiguously up to the family level. Amplicon sequences may also be assigned as simply k Fungi. In some cases these amplicons might truly be from fungal taxa that are just not well-represented in a given curated fungal database, but metazoan amplicons, for example, can also be assigned to this “Unknown Fungus” taxonomy. If metazoan outgroup sequences are present in a database, these unknown metazoan amplicons should theoretically be more similar to those outgroups and be assigned to k Metazoa.

Here, we compare taxonomic assignments from 15 published fungal meta-amplicon studies (Table 1) using both the UNITE_Fungi (Abarenkov et al., 2022) and UNITE_All (Abarenkov et al., 2022a) databases to test the effects of including outgroups in taxonomic assignments. We examine agreement between assignments at each taxonomic level and discuss potential implications for ecological inferences. We hypothesized that the UNITE_All database (containing fungi and other eukaryote sequences) would help to detect and remove non-fungal reads that might simply be assigned as “unknown fungus” when using the UNITE_Fungi database. Our results suggest that many fungal meta-amplicon studies may be unwittingly including non-fungi in their estimates of diversity, and that the inclusion of outgroups in taxonomic databases is essential for accurate ecological measures in fungal meta-amplicon studies.

**Table 1:** The publically available data sets used in this study.

| Host | Habitat | Taxa | Shannon Diversity | SRA Accession |
| --- | --- | --- | --- | --- |
| Soil | Temperate | 1177 | 3.11 | PRJNA480528 |
| Soil | Tropical | 2725 | 4.19 | PRJNA379981 |
| Soil | Desert | 1049 | 2.91 | PRJNA659159 |
| Soil | Tropical | 10823 | 6.06 | PRJNA679063 |
| Soil | Tropical | 203 | 1.62 | PRJNA316729 |
| Plant | Tropical | 118 | 1.73 | PRJNA474551 |
| Plant | Marine | 591 | 3.72 | PRJNA515720 |
| Plant | Tropical | 4911 | 3.95 | PRJNA355011 |
| Plant | Marine | 178 | 4.03 | PRJNA667465 |
| Animal | Laboratory | 4085 | 3.85 | PRJNA597168 |
| Animal | Tropical | 1479 | 1.87 | PRJNA634617 |
| Animal | Laboratory | 1175 | 1.97 | PRJNA512838 |
| Aerobiota | Coastal | 5152 | 5.31 | PRJNA736519 |
| Aerobiota | Coastal | 3862 | 3.94 | PRJNA736543 |
| Aquatic | Freshwater | 1050 | 2.68 | PRJNA436133 |

## MATERIALS AND METHODS

### Overview

We selected 15 fungal meta-amplicon studies with publicly available raw ITS amplicons. Raw data for each study was downloaded and processed into ASV tables. All ASV tables were combined into a single table and taxonomy was assigned to ASVs using both a fungal-only database (UNITE_Fungi) and a database containing fungi and eukaryotic outgroups (UNITE_All). Taxonomic assignments were made using both the RDP Classifier and by selecting the top BLAST hit from a standalone blastn call against both databases. All analyses were performed in R (version 4.2.0).

### Data aquisition

We searched the literature for studies that reported fungal mycobiome data from either ITS1 or ITS2 amplicons. Studies were selected for inclusion based on availability of raw data in the Sequence Read Archive (SRA) and unambiguous metadata (Table 1). Sequences were downloaded directly from the SRA using the fasterq-dump module from the sra-toolkit (version 2.10.9; https://github.com/ncbi/sra-tools). Metadata from each study was downloaded directly from SRA. Scripts for downloading data are included in Supporting Info.

### Data processing and taxonomic assignment

Raw data from each study was processed as follows: First, the ITS regions were extracted using *itsxpress* (Rivers et al., 2018). Next, extracted forward reads were subjected to quality filtration using the *dada2* R package (Callahan et al., 2016) where sequences with uncalled bases were removed and sequences were truncated when quality scores dropped below a phred score of 25. Quality filtered reads were denoised with the DADA2 algorithm and processed into ASV tables for each study. Each study was combined with metadata from the SRA and placed into a phyloseq object using the *phyloseq* R package (McMurdie & Holmes, 2013). All phyloseq objects were combined into a single object and taxonomy was assigned using the assign_taxonomy() function from *dada2* using both the UNITE_Fungi and UNITE_All databases. ASV sequences were also exported for assignment against those two databases using the blastn tool from BLAST (version 2.13.0). The top BLAST hit, based on e-value and coverage, was selected as the “BLAST taxonomy.” These various taxonomic assignments were added to phyloseq objects for all downstream analyses.

### Analyses

Taxonomic assignments were compared at each taxonomic level to examine agreement between the UNITE_Fungi and UNITE_All databases. Agreement was indicated as exact taxonomic match at a given level for each ASV. After agreement values were determined, we removed any taxa not assigned to Kingdom Fungi for estimating fungal diversity. All ASVs identified as non-fungal by both the BLAST and RDP Classifier methods against the UNITE_All database were investigated under the corresponding ASV assignments by the UNITE_Fungi database. All analysis code can be found in the GitHub repository associated with this manuscript (https://github.com/gzahn/Fungal_Database_Comparison).

## RESULTS

Out of 40,232 unique ASVs recovered from all the studies, 16,106 were determined to be of non-fungal origin based on RDP Classifier and BLAST assignments against the UNITE_All database. Of these, taxonomic assignment against the UNITE_Fungi database falsely identified 16,075 of them as kingdom fungi (See Supporting Info). The majority of non-fungal ASVs that were falsely identified as “unknown fungus” belonged to either unknown or metazoan kingdoms (Figure 1). Since none of the studies in question described any methods for removing non-fungi, these ASVs presumably remained in the studies in question, contributing to “fungal” diversity measures.

**Figure 1:**
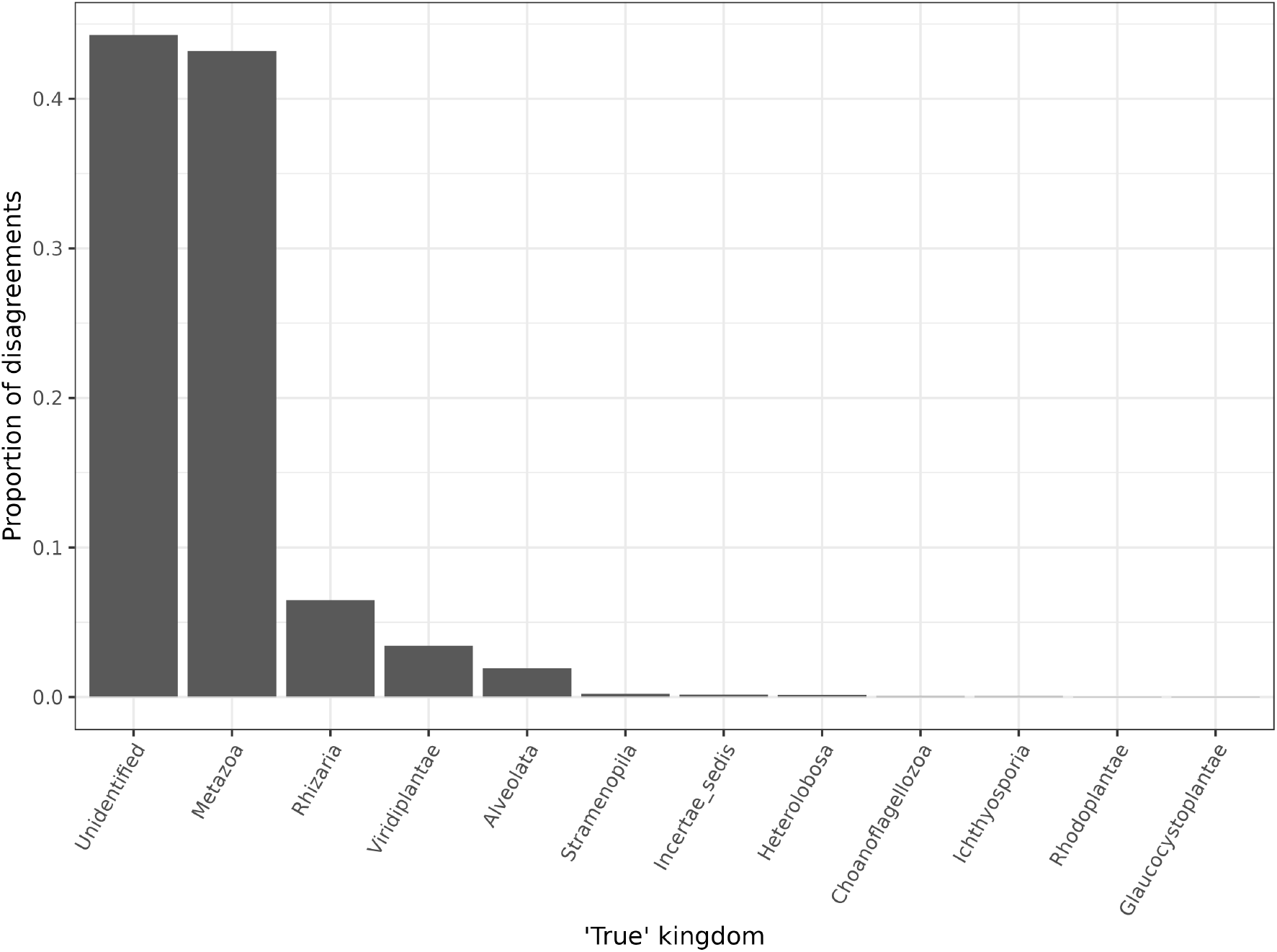
The proportion of kingdoms represented by non-fungal ASVs that were falsely identified as “*Fungus sp*.” by the UNITE_Fungi database.

Taxonomic agreement between the two databases varied between studies and between major habitats. Disagreements were largely driven by disparities at the kingdom level, but were more pronounced at more specific taxonomic levels (Figure 2). ASVs from aerial and soil habitats showed fewer falsely assigned fungi while ASVs from plant and animal hosts had proportionally more falsely assigned fungi. Limited replication reduced our statistical power so that these patterns were not statistically significant, however.

**Figure 2:**
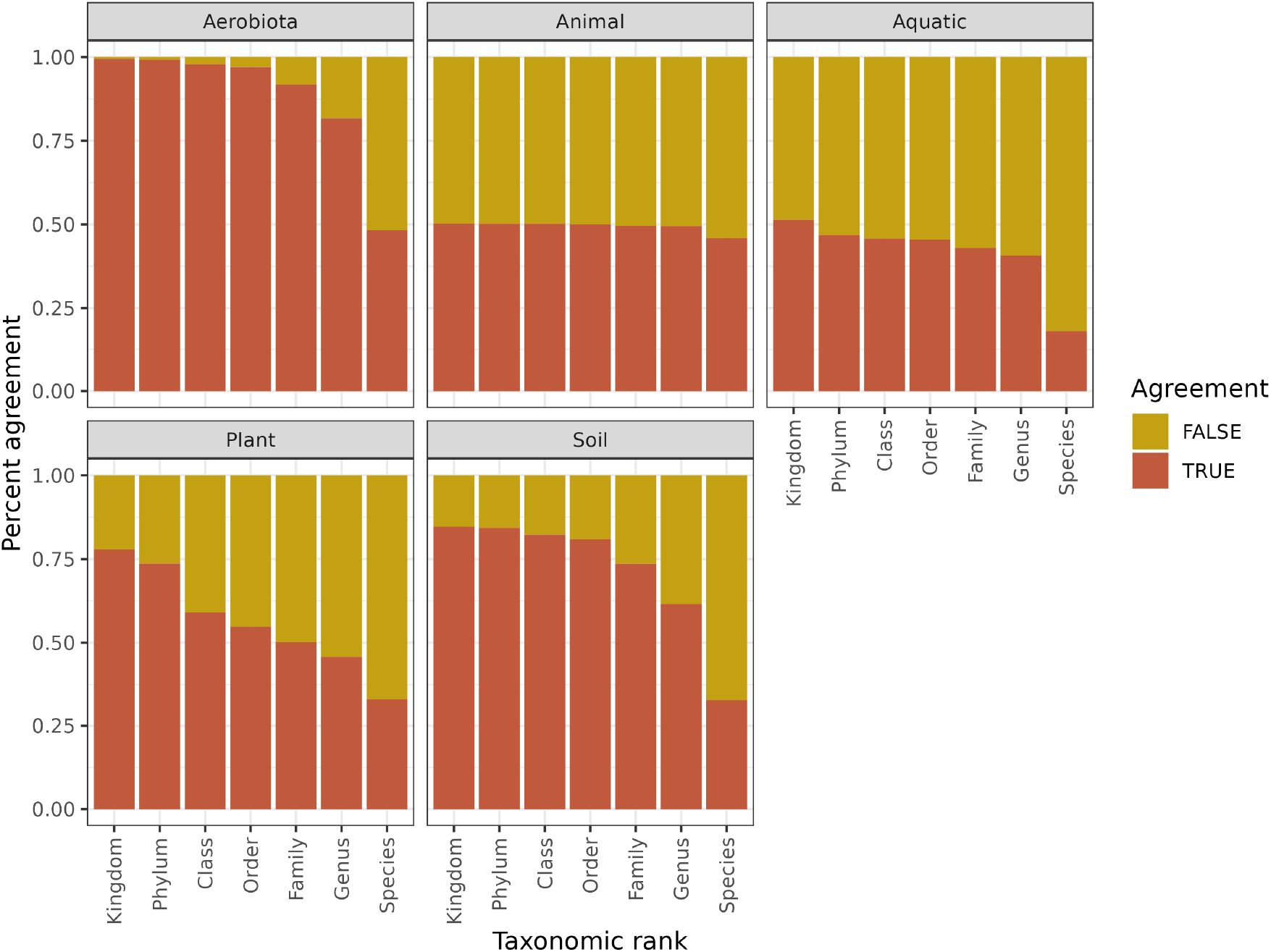
The proportional agreement between UNITE_Fungi and UNITE_All databases shown at the ecosystem level.

Alpha diversity (richness) was affected by these falsely assigned “fungal” ASVs (Figure 3). Comparing the fungal richness between database methods showed artificial inflation of “fungal” diversity when ASVs were assigned taxonomy using the UNITE_Fungi database. The vast majority of these assignments were simply “*Fungi sp*.” so it is unlikely to have affected any lineage-specific conclusions reached in the associated studies. However, it seems plausible that these and other studies that use the UNITE_Fungi database without outgroups will show inflated alpha diversity measures. Some ecosystems may be more affected by this issue than others.

**Figure 3:**
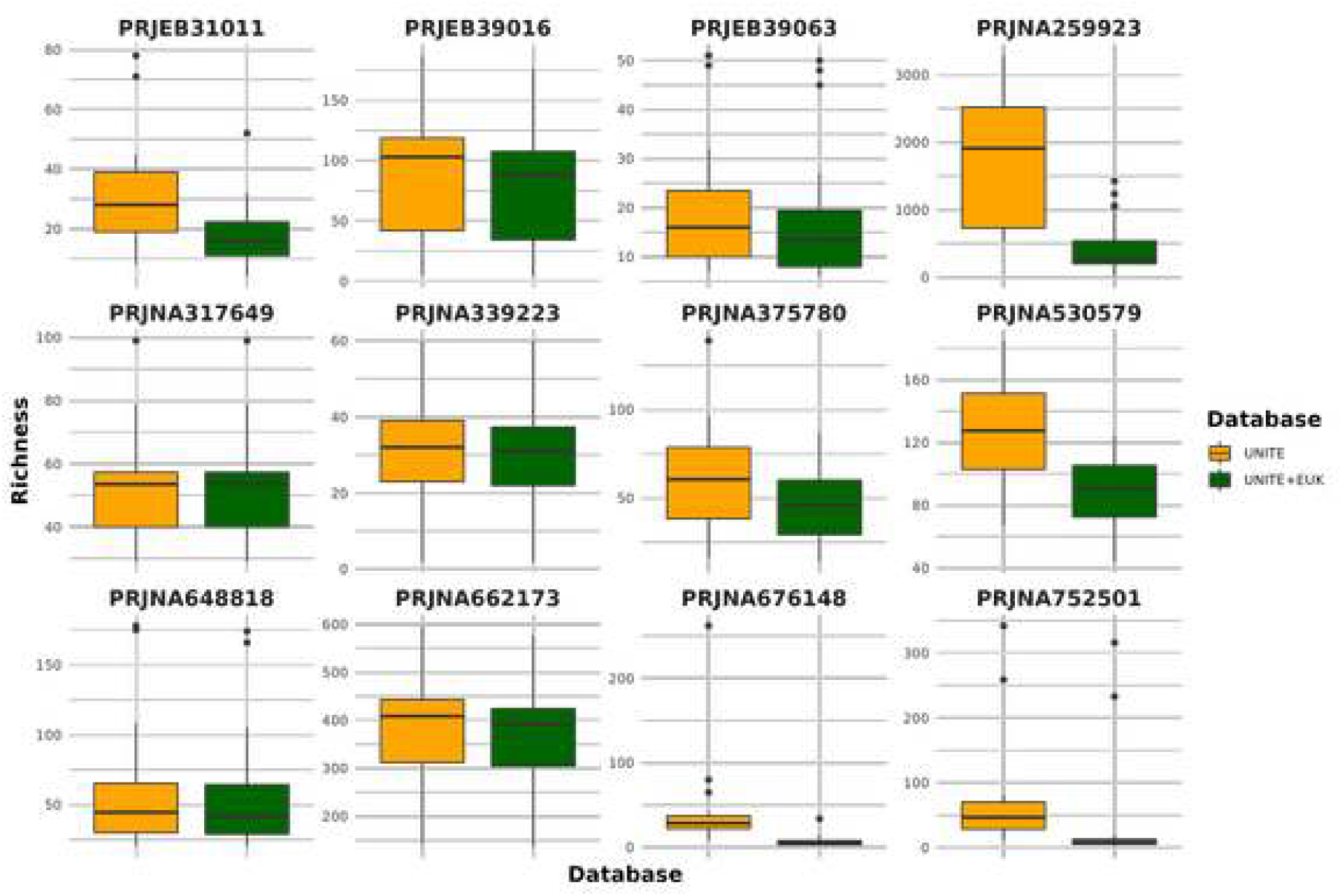
Alpha diversity of each study after removing non-fungi; Falsely assigned fungi were not removed and inflated alpha diversity measures. Only 12 studies shown.

## DISCUSSION

Alpha diversity measures are a common tool for comparing samples within and between studies. If a goal of a study is to examine fungal diversity, care must be taken to remove non-fungal sequences. This depends on reliable taxonomic assignments of those sequences. By assigning taxonomy with a database that *only* includes fungi, the chances of non-fungal sequences being assigned as some unspecified “*Fungus sp*” are dramatically increased. In this analysis of 15 published studies, we found that 99.8% of all non-fungal sequences were assigned to “*Fungus sp*.” when using the UNITE_Fungi database.

The ability of common fungal primers to co-amplify non-target templates has been discussed before (Bellemain et al., 2010; Ihrmark et al., 2012; Tedersoo & Anslan, 2019). While *in silico* tests may help to quantify the danger of this occurring for specific primer sets and specific conditions, even the most careful PCR protocol is not a homogenous reaction. Temperature variations and other conditions can lead to non-specific binding. For studies of the fungal mycobiome where host or environmental DNA is present, there will always be a risk that other eukaryotic taxa are amplified.

Our results highlight that this has occurred and gone undetected in the published literature (Figure 4), but that it is not an insurmountable problem. It can be mitigated by simply including outgroups in the database when taxonomy is assigned and then by removing taxa not unambiguously assigned to the fungal kingdom. We strongly suggest that future studies of this nature use the UNITE_All database for assigning taxonomy.

**Figure 4:**
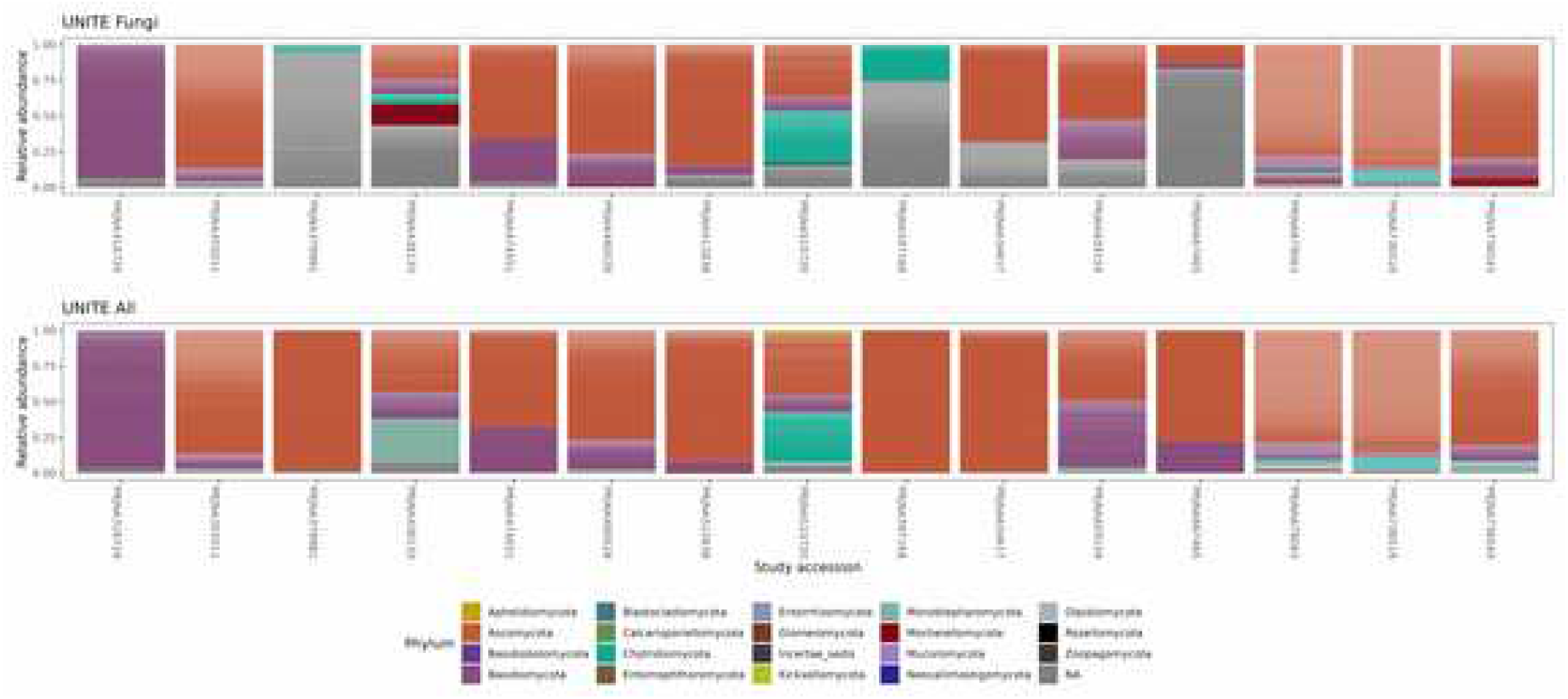
Phylum relative abundance after removing non-fungi from each taxonomic assignment method. The bulk of “Fungus sp.” that were assigned using UNITE_Fungi had no phylum-level assignment (Gray fill in this plot).

## Supporting information

Fasta file of all ASV sequences analyzed

## DATA AVAILABILITY STATEMENT

All raw sequenced used in this study come from previously published work and are publicly available on the Sequence Read Archive under their listed accessions (Table 1). All analysis code is publicly available at https://github.com/gzahn/Fungal_Database_Comparison

